# Genome structural variation in natural *Caenorhabditis elegans* populations

**DOI:** 10.1101/2025.01.07.631790

**Authors:** Kyle J. Lesack, James D. Wasmuth

**Affiliations:** Faculty of Veterinary Medicine, University of Calgary, Calgary, Alberta, Canada

**Keywords:** structural variation, *Caenorhabditis elegans*, environmental adaptation

## Abstract

Structural variation, involving large alterations in chromosome structure, drives genetic diversification and the emergence of new phenotypes. These changes are widespread in natural populations and play an important role in adaptation and speciation. For many species, research has been limited to laboratory adapted strains and experimental evolution, which may not reflect the diversity of structural variants in the wild. Furthermore, technological limitations have proved to be a major barrier to accurate and comprehensive variant calling. In this study, we used PacBio sequencing data from 14 wild *Caenorhabditis elegans* strains to characterize structural variants in a natural population. With long-reads, we overcame limitations associated with short-read approaches and leveraged population-level data to further refine the accuracy of variant calls. We found that large, rapidly evolving gene families, such as GPCR, F-Box, and C-Type lectin, were prominent among the variants predicted to have phenotypic consequences. These results shed light on the significant role of structural variants in the evolution of *Caenorhabditis elegans* in its wild habitats and the limitations of treating N2 as the reference wild-type.

## Introduction

Structural variants (SVs) are large-scale alterations in a genome—such as deletions, duplications, and inversions—that affect gene content, order, or dosage. These variants can have profound phenotypic effects by disrupting genes, altering gene dosage, or rearranging regulatory elements. While often deleterious, SVs can also drive adaptive evolution through mechanisms such as gene duplication followed by neofunctionalization or chromosomal inversions that suppress recombination and maintain beneficial allele combinations (Faria et al., 2019; Katju and Bergthorsson, 2013; Kirkpatrick, 2010).

Despite their importance, SVs remain poorly characterized in most species, especially at the population level. This is partly due to the technical limitations of short-read DNA sequencing, which struggles to resolve large or complex variants, particularly in repetitive regions (Koboldt, 2020; Lesack et al., 2022). Long-read sequencing has substantially improved the resolution of SVs and providing more reliable alignments (Smolka et al., 2024).

In the nematode *Caenorhabditis elegans*, much of our knowledge about SVs comes from laboratory strains and mutation accumulation experiments. These studies have revealed a high rate of spontaneous duplications and the formation of complex alleles (Farslow et al., 2015; Konrad et al., 2018). For example, a recombinant inbred line showed increased fitness due to a partial duplication and inversion of five different regions of *rcan-1*, a gene which regulates thermotactic behavior (Zhao et al., 2020). However, laboratory strains, such as N2, carry numerous derived alleles and pseudogenised genes, limiting their relevance for understanding variation in natural populations (Andersen and Rockman, 2022; Sterken et al., 2015).

Comparative studies between N2 and wild isolates, particularly the Hawaiian strain CB4856, have revealed extensive structural divergence (Lee et al., 2021; Vergara et al., 2014). In 12 natural strains, more than 5% of the N2 genome is affected by SVs, with CB4856 showing over 79,000 insertion/deletion events, more than 800 unaligned segments, and 61 hyper-divergent regions (Lee et al., 2021; Thompson et al., 2015).

The *Caenorhabditis elegans* Natural Diversity Resource (CeNDR) has expanded access to genomic data from wild strains (Cook et al., 2017). This collection has facilitated the discovery of genetic loci underlying xenobiotic metabolism and anthelmintic drug resistance (Hahnel et al., 2018; Zdraljevic et al., 2019).

In this study, we used PacBio long-read sequencing data from 14 wild *C. elegans* strains to generate a high-confidence, population-scale map of SVs. We identified thousands of deletions, duplications, and inversions—many affecting genes involved in environmental response, stress, and immune function.

## Results and Discussion

We identified 14,373 deletions, 3,149 duplications, and 1,748 inversions in 14 wild *C. elegans* strains (Table S1). SVs were unevenly distributed across the genome (Figure S1). Deletions and duplications were enriched at the autosomal arms, whereas inversions were more evenly distributed. These patterns are consistent with previous findings; SVs preferentially occur in low gene density, high recombination regions (Cutter et al., 2009). Chromosome arms may promote SV formation through recombination-based mechanisms while protecting essential genes located in denser chromosome centres (Carvalho and Lupski, 2016). Among all SVs, 64% were singletons—variants unique to a single strain—suggesting high inter-strain diversity (Table S2). Hawaiian strains had the highest number and proportion of singleton variants while Western European strains had the fewest (Table 1). The high ancestral diversity in Hawaiian strains is known, with reduced diversity in non-Hawaiian lineages due to selective sweeps (Crombie et al., 2019). Conversely, 25 variants were found in all 14 strains: 15 deletions, two duplications, and eight inversions (Table S2; Figures S2, S3, S4).

**Table 1.** Singleton variants among individual *C. elegans* strains.

| Strain | Region | Deletions |  |  | Duplications |  |  | Inversions |  |  |
| --- | --- | --- | --- | --- | --- | --- | --- | --- | --- | --- |
|  |  | Total | Singleton | Singleton Proportion | Total | Singleton | Singleton Proportion | Total | Singleton | Singleton Proportion |
| DL238 | Hawaii | 2343 | 804 | 0.34 | 485 | 209 | 0.43 | 333 | 95 | 0.29 |
| ECA396 | Hawaii | 3412 | 1643 | 0.48 | 519 | 300 | 0.58 | 399 | 140 | 0.35 |
| QX1794 | Hawaii | 2363 | 756 | 0.32 | 459 | 186 | 0.41 | 311 | 74 | 0.24 |
| XZ1516 | Hawaii | 4007 | 2440 | 0.61 | 443 | 266 | 0.60 | 519 | 301 | 0.58 |
| ECA36 | New Zealand | 3522 | 1671 | 0.47 | 586 | 308 | 0.53 | 398 | 147 | 0.37 |
| EG4725 | Western Europe | 1913 | 290 | 0.15 | 452 | 94 | 0.21 | 280 | 42 | 0.15 |
| JU1400 | Western Europe | 1692 | 225 | 0.13 | 417 | 120 | 0.29 | 240 | 34 | 0.14 |
| JU2526 | Western Europe | 2257 | 452 | 0.20 | 442 | 96 | 0.22 | 309 | 55 | 0.18 |
| JU2600 | Western Europe | 1519 | 290 | 0.19 | 385 | 87 | 0.23 | 238 | 39 | 0.16 |
| JU310 | Western Europe | 1366 | 170 | 0.12 | 385 | 72 | 0.19 | 217 | 16 | 0.07 |
| MY2147 | Western Europe | 1279 | 217 | 0.17 | 364 | 85 | 0.23 | 199 | 29 | 0.15 |
| MY2693 | Western Europe | 1251 | 97 | 0.08 | 385 | 84 | 0.22 | 258 | 35 | 0.14 |
| NIC2 | Western Europe | 457 | 80 | 0.18 | 248 | 77 | 0.31 | 150 | 12 | 0.08 |
| NIC526 | Western Europe | 1544 | 179 | 0.12 | 402 | 78 | 0.19 | 257 | 22 | 0.09 |

To prioritise variants with likely functional consequences, we focused on high-impact SVs. This subset included 1,485 deletions, 214 duplications, and three inversions, collectively affecting 1,258 genes (Figure 1; Tables S3, S4, S5). Deletions were enriched for splicing disruptions, transcript ablations, and frameshift variants (Table 2). Duplications primarily resulted in transcript amplifications, and all three high impact inversions were classified as stop/loss mutations. Functional enrichment revealed strong overrepresentation in categories related to environmental stress, including GPCRs, F-box proteins, C-type lectins (CLECs), and detoxification enzymes (Figure 2; Table S6). Many of these genes were in hyper-variable regions, which have been shown to maintain ancestral variation through balancing selection (Lee et al., 2021).

**Figure 1.**
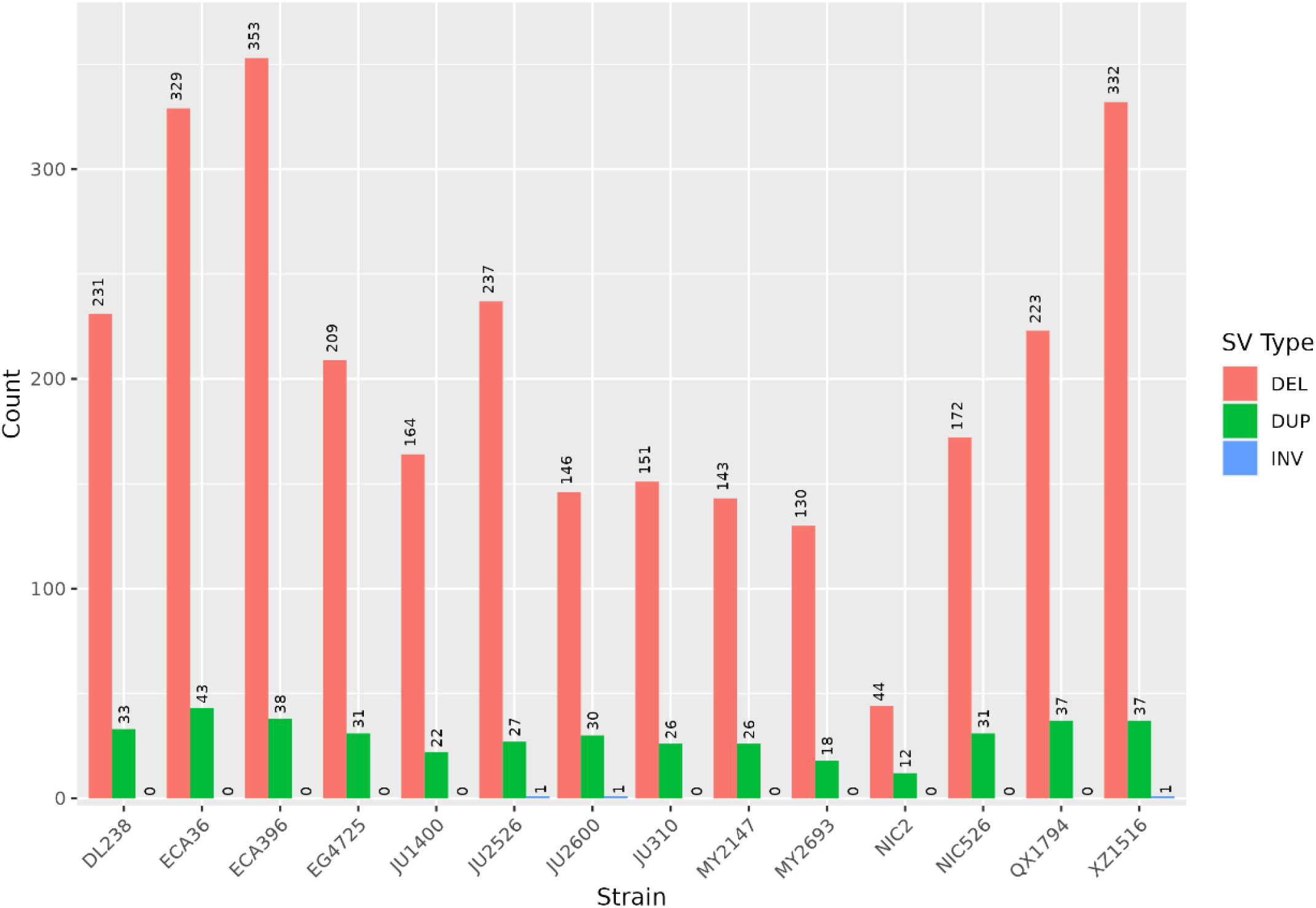
Number of structural variants determined to have high functional impact.

**Figure 2.**
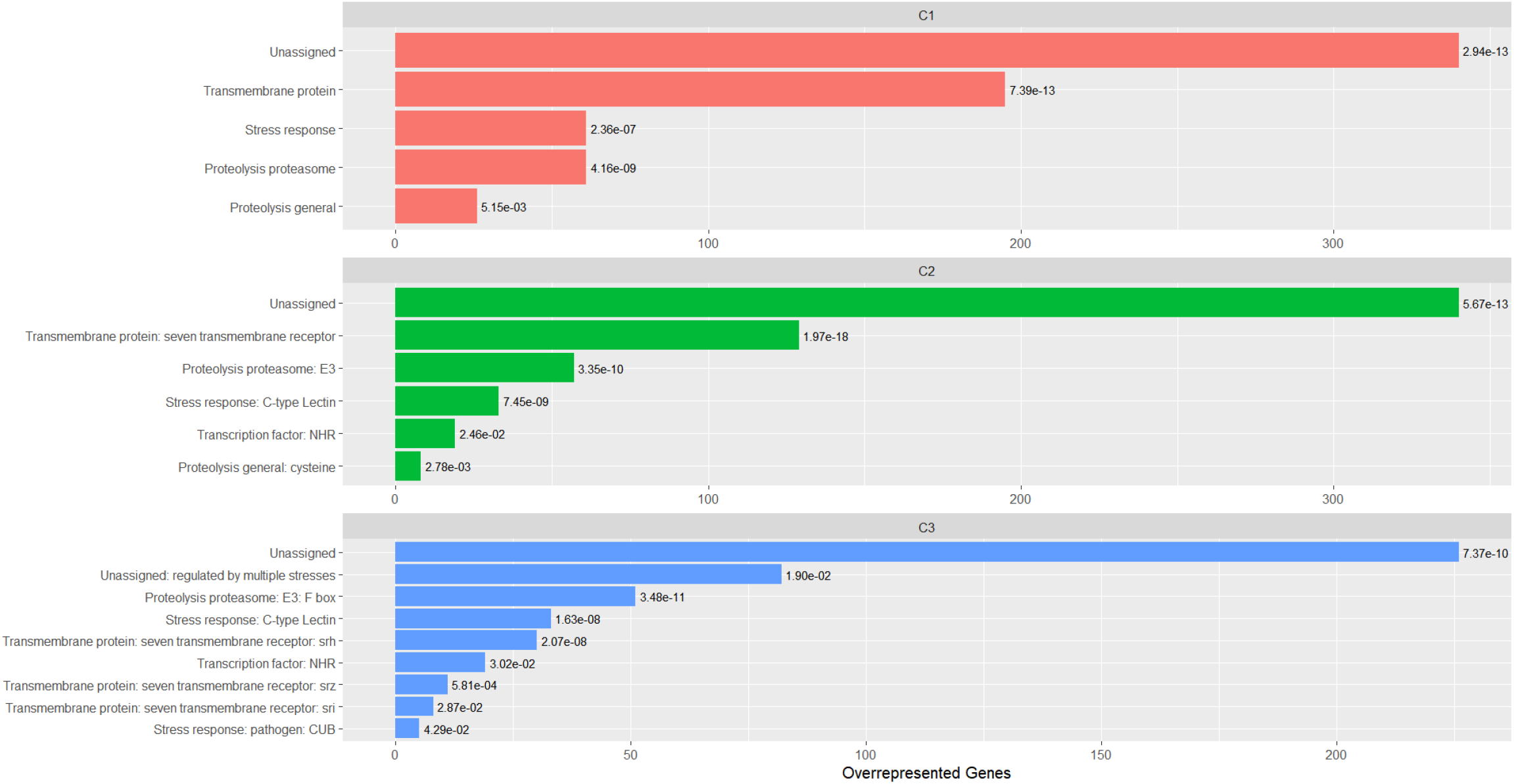
Statistically significant overrepresented WormCat terms for putative high impact SVs. The red, green, and blue colours represent the categories for the C1, C2, and C3 levels, respectively.

**Table 2.** Predicted Consequences of High Severity Structural Variants. **‡** Consequence descriptions downloaded on May 22, 2023 from: https://useast.ensembl.org/info/genome/variation/prediction/predicted_data.html

| SV Type | Term | Description‡ | Count |
| --- | --- | --- | --- |
| Deletion<br>(n = 1,485) | splice_donor_variant | A splice variant that changes the 2-base region at the 5' end of an intron | 510 |
|  | splice_acceptor_variant | A splice variant that changes the 2-base region at the 3' end of an intron | 503 |
|  | transcript_ablation | A feature ablation whereby the deleted region includes a transcript feature | 240 |
|  | frameshift_variant | A sequence variant which causes a disruption of the translational reading frame, because the number of nucleotides inserted or deleted is not a multiple of three | 168 |
|  | stop_lost | A sequence variant where at least one base of the terminator codon (stop) is changed, resulting in an elongated transcript | 89 |
|  | start_lost | A codon variant that changes at least one base of the canonical start codon | 16 |
|  | stop_gained | A sequence variant whereby at least one base of a codon is changed, resulting in a premature stop codon, leading to a shortened transcript | 2 |
| Duplication<br>(n = 241) | transcript_amplification | A feature amplification of a region containing a transcript | 240 |
|  | start_lost | A codon variant that changes at least one base of the canonical start codon | 1 |
| Inversion<br>(n=3) | start_lost | A codon variant that changes at least one base of the canonical start codon | 3 |

### GPCRs and sensory behaviour

Seven-transmembrane G-protein-coupled receptors (GPCRs), involved in chemosensation and behaviour, were among the most affected gene families. Of the 195 genes in the “Transmembrane protein” category, 120 were GPCRs. An additional GPCR, *srh-208*, was categorized under the “Proteolysis proteasome” category. The SRH family was enriched, with 31 genes, which is consistent with previous reports of rapid expansion and diversification in *C. elegans* (Thomas and Robertson, 2008; Vidal et al., 2018). A duplication of *srh-234* was present in two strains. The expression of *srh-234* is regulated by the chemosensation of food (Gruner et al., 2014). The deletion of *str-208* was noteworthy given that this gene is mplicated in erratic movement (Daigle et al., 2022). While the loss of this gene may affect locomotion, it is not yet known whether the associated structural variants confer an adaptive benefit, impose a cost, or are selectively neutral.

### F-box genes and immune adaptation

SVs affected 87 proteolysis-related genes, including 44 F-box genes. The F-box gene family is considerably expanded in *C. elegans* relative to other *Caenorhabditis* species with several members under positive selection in the non-Hawaiian strains (Ma et al., 2021). Additional immune-related SVs may reflect adaptations to local microbial environments. For example, *hecw-1*, deleted in ECA-396 is an E3 ubiquitin ligase that modulate pathogen avoidance (Chang et al., 2011).

### C-type lectins and pathogen response

Thirty-one C-type lectin (CLEC) genes were affected by SVs. CLECs are a rapidly evolving gene family involved in innate immunity and pathogen recognition (Pees et al., 2016). Five CLECs were affected by deletions—*clec-2, clec-10, clec-122, clec-143*, and *clec-174*—all known to be modulated by bacterial pathogens (Pan et al., 2021; Sahu et al., 2012; White and Herman, 2018). An additional CLEC, *clec-189*, was predicted as being deleted in EG4725 and JU2526, partially deleted in QX1794 and XZ1516, and duplicated in EG4725 and JU2526. Although it awaits functional annotation it may represent a novel immune effector (Pan et al., 2021). The prediction of complete deletions and duplications spanning a single gene within the same strains indicates that even long-read SV callers require further improvements.

### Detoxification and xenobiotic response

Several families of detoxification genes were enriched in SVs, including cytochrome P450s (CYPs), glutathione S-transferases (GSTs), UDP-glucuronosyltransferases (UGTs), and nuclear hormone receptors (NHRs). Two caffeine inducible CYPs, *cyp-33e2* and *cyp-35a4* harbored deletions; the latter gene was also duplicated in QX1794 and is associated with the regulation of fat storage (Aarnio et al., 2011; Min et al., 2015). Other deletion-affected CYP genes included: *cyp-33c5* and *cyp-13a5*, which are involved in cadmium detoxification; and *cyp-33d3*, which is inducible by ethidium bromide and low pH (Cong et al., 2020; Cui et al., 2007). A deletion was also identified in *ugt-14*, which is upregulated in response to benzimidazole*-*class anthelmintics (Stasiuk et al., 2019).

### Evolutionary and Functional Implications

Our results highlight the utility of long-read sequencing for characterizing SVs at a population scale. The high diversity of SVs in wild *C. elegans* populations—particularly those in gene families related to sensory perception, immunity, and detoxification—emphasizes the limitations of relying solely on N2 as a wild-type reference (Andersen and Rockman, 2022; Sterken et al., 2015). Many of the identified SVs may confer adaptive advantages, particularly in geographically and microbially diverse environments.

Importantly, many SVs overlap with hyper-divergent regions—genomic hotspots for adaptive diversity—shown in *C. elegans* and the parasitic *Heligmosomoides polygyrus* to contain genes involved in sensory perception and stress response (Lee et al., 2021; Stevens et al., 2023).

Going forward, hybrid approaches that integrate long- and short-read sequencing and transcriptomic data from wild isolates will help resolve the regulatory and phenotypic consequences of SVs (Zhang et al., 2022). In particular, eQTL mapping that includes SVs could offer insights into complex traits not explained by single-nucleotide variation alone (Boyling et al., 2022). Additionally, functional studies targeting SVs genes currently categorized as “Unassigned” may reveal previously uncharacterized molecular mechanisms.

## Materials and Methods

### Strains

We analysed PacBio sequencing data from 14 *C. elegans* strains (DL238, ECA36, ECA396, EG4725, JU1400, JU2526, JU2600, JU310, MY2147, MY2693, NIC2, NIC526, QX1794, XZ1516) sourced from CeNDR and collected from Hawaii, New Zealand, and Western Europe (Figure S5).

### Sequence alignment and structural variant calling

Reads were aligned to the N2 reference genome using NGMLR (v0.2.7), and alignments were sorted using Picard (v2.27.5) (Broad Institute, 2025; Sedlazeck et al., 2018). SVs were called using Sniffles2 (v2.0.7), filtered to include only deletions, duplications, and inversions ≥ 100bp (Smolka et al., 2024). SV calls were refined using Iris (v1.0.04) for improved breakpoint resolution and merged across strains using Jasmine (v1.1.5) and Iris (v1.0.4) (Kirsche et al., 2023).

### SV annotation and functional profiling

We used Ensembl’s Variant Effect Predictor (v.109.3) to predict the functional consequences of each SV (McLaren et al., 2016). SVs classified as “high impact” were selected for functional enrichment using easyGSEA (v1.3.1) and annotated using WormCat (v2) (Cheng et al., 2021; Holdorf et al., 2020). Genes were assigned to statistically significant terms across WormCat’s hierarchical levels (FDR ≤ 0.05), though no single level was exclusively used due to trade-offs in resolution and coverage. Gene family membership was assigned based on the annotations in the WormBase database.

### Data presentation

SV densities along chromosomes were plotted using karyoploteR (v1.26.0) (Gel and Serra, 2017). SV allele frequencies were visualized UpSetPlot (v0.8.0) (Lex et al., 2014). Only the largest intersections—25 duplications, 20 for deletions and inversions—were displayed. The remaining plots were created using ggplot2 (v.3.4.2) (Wickham, 2016).

## Data availability

The code, and Snakemake pipeline, used in this project is freely available: https://github.com/kyleLesack/sv_analysis_cendr_pacbio

## Acknowledgments

We thank Dr. Erik Andersen and members of the Andersen lab (Johns Hopkins University) for discussions on using CeNDR datasets. We also acknowledge the high-performance computing resources by the Faculty of Veterinary Medicine and Research Computing at the University of Calgary.

## Funding

This work was funded through grants to JDW from the Natural Sciences and Engineering Research Council of Canada (NSERC; #04589-2020).

## Notes

### Competing Interest Statement

The authors have declared no competing interest.

### Summary of Updates

Small changes in the results and associated figures.

