## Supplemental tables and figures for "Genome structural variation in natural *Caenorhabditis elegans* populations"

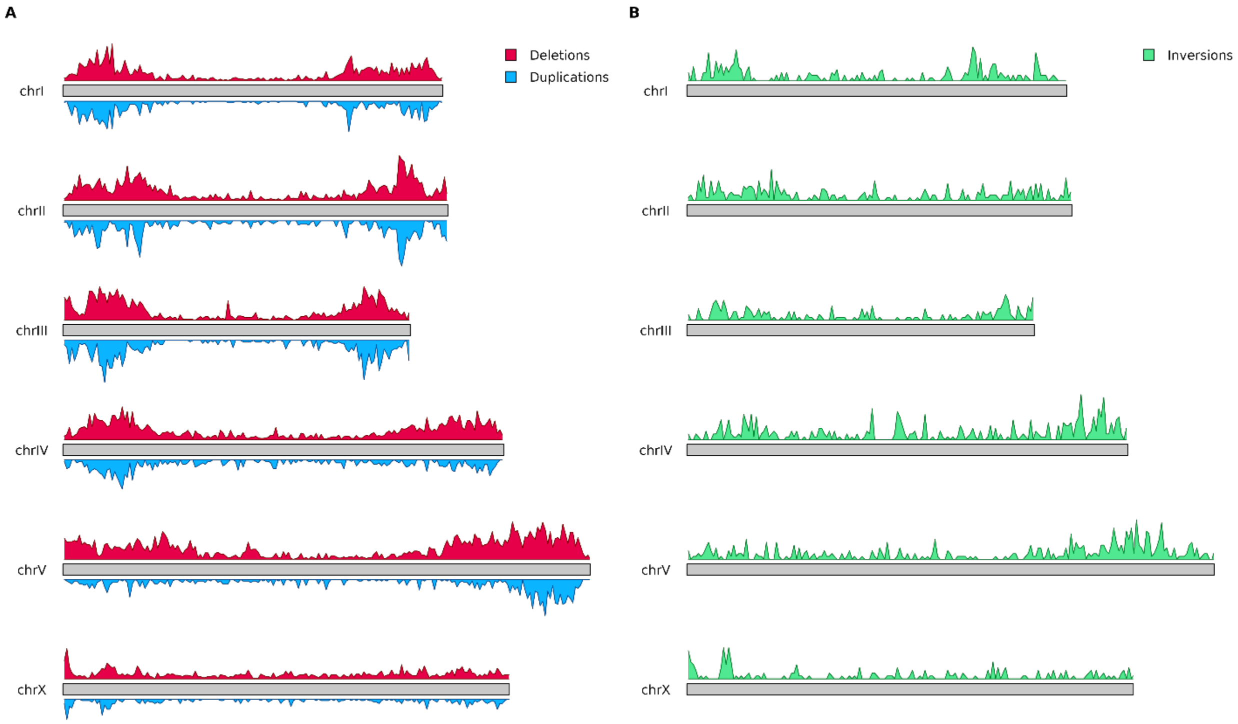


**Figure S1:** Distribution of SVs along chromosomes. (A) Deletions and duplications are shown in blue and red, respectively. (B) Inversions are shown in green.

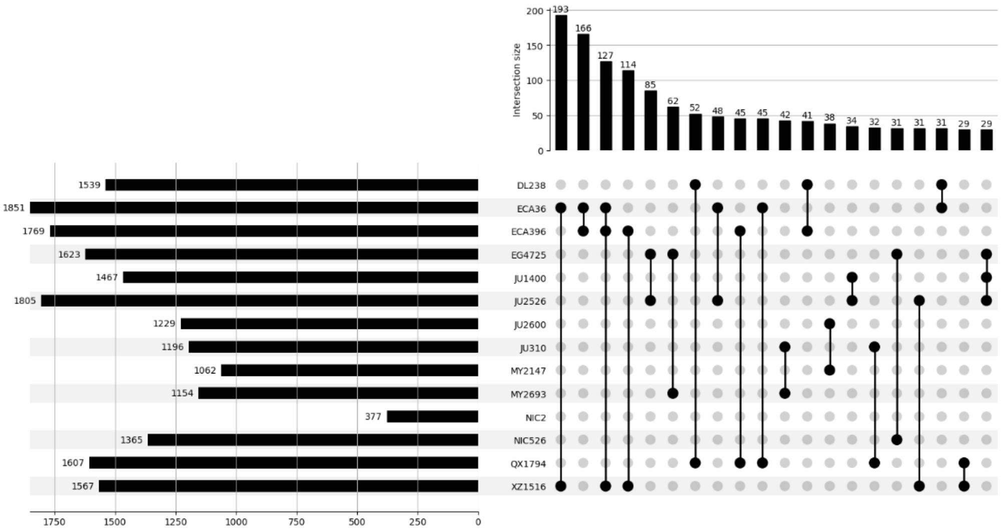


**Table S2:** Distribution of deletion allele frequencies in wild *C. elegans* strains. The count totals only include variants present in at least two strains. The plot is limited to the 20 largest sets.

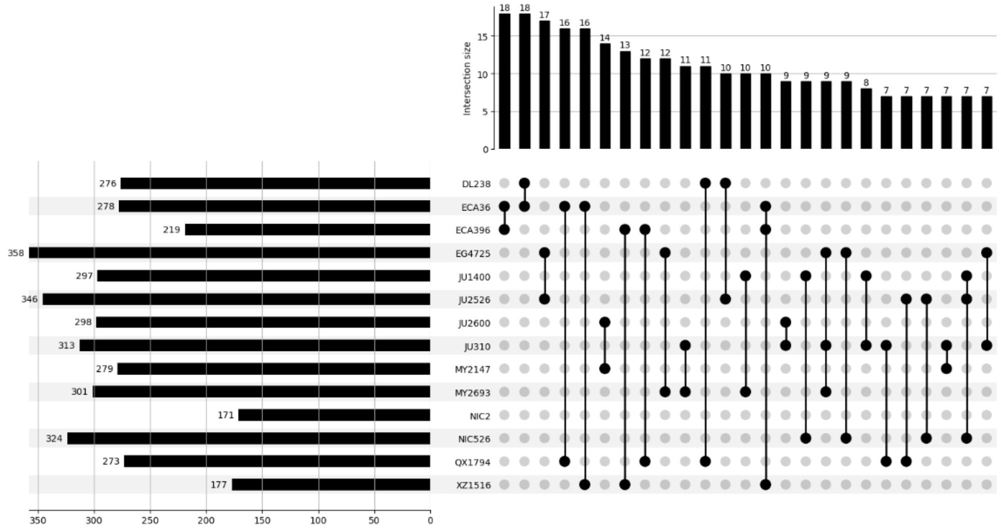


**Figure S3:** Distribution of duplications allele frequencies in wild *C. elegans* strains. The count totals only include variants present in at least two strains. The plot is limited to the 25 largest sets.

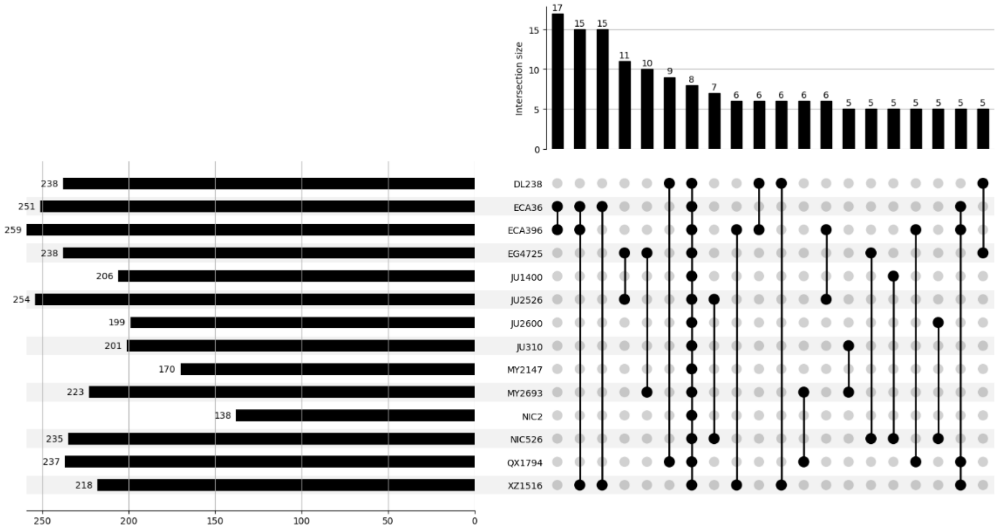


**Figure S4:** Distribution of inversion allele frequencies in wild *C. elegans* strains. The count totals only include variants present in at least two strains. The plot is limited to the 20 largest sets.

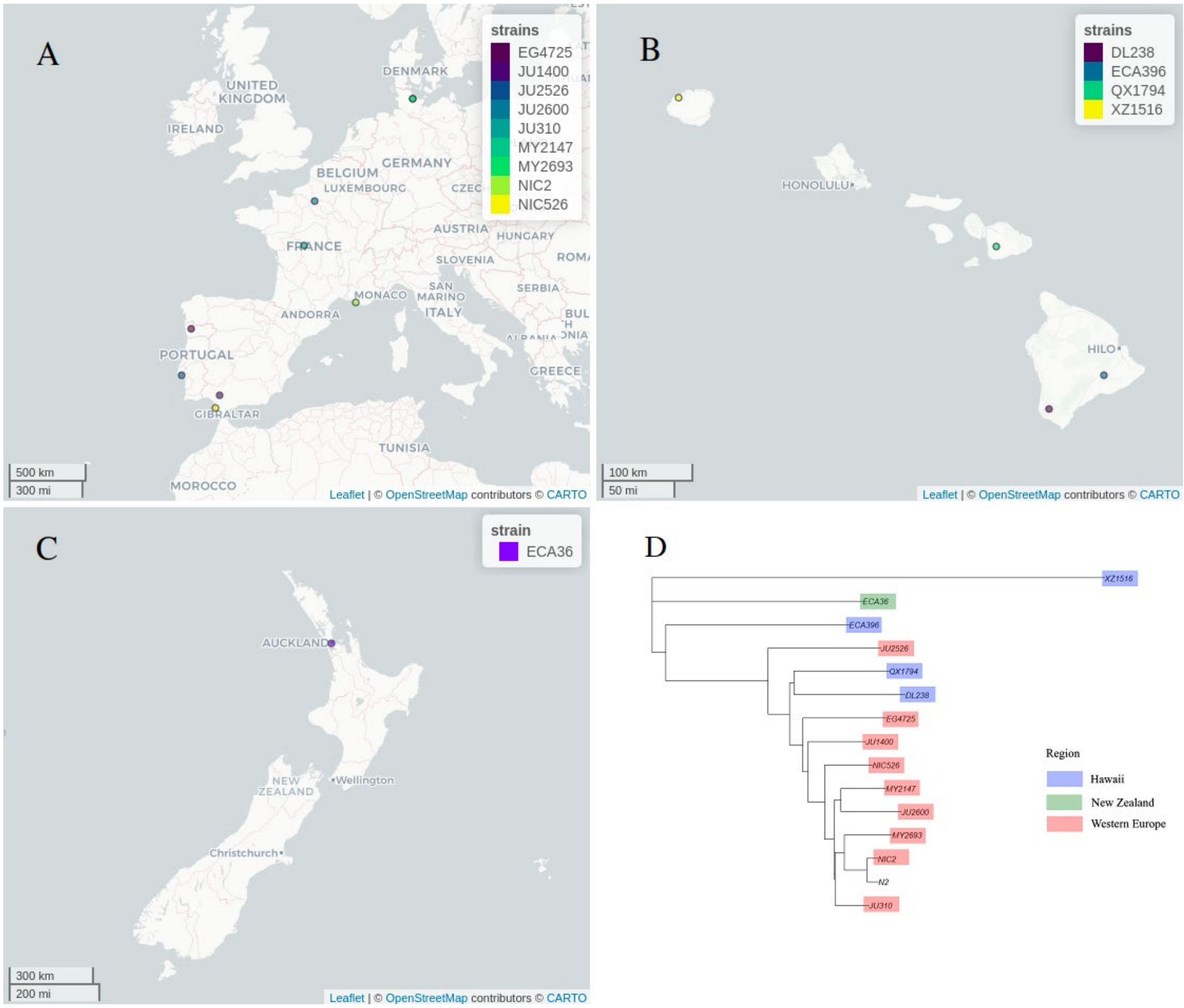


**Figure S5:** CeNDR sampling. (A) Nine strains were collected from Western Europe. (B) Four strains were collected from Hawaii. (C) One strain was collected from New Zealand. (D) Phylogenetic tree for the 14 strains (adapted from the CeNDR species tree available at https://www.elegansvariation.org/data/release/latest).

**Table S1:** Summary of structural variants

| Variant Type | Count | Mean Size ±SD (bp) | Minimum Size (bp) | Maximum Size (bp) |
|---|---|---|---|---|
| Deletion | 14,373 | 705 ±2,459 | 100 | 97,753 |
| Duplication | 3,149 | 1,523±4,163 | 103 | 75,314 |
| Inversion | 1,748 | 3,822 ±8,684 | 100 | 96,813 |

**Table S2:** Presence of structural variants in multiple strains

| **Number of strains containing variant†** | **Deletions** | **Duplications** | **Inversions** |
|---|---|---|---|
| 1 | 9314 | 2062 | 1041 |
| 2 | 1897 | 456 | 245 |
| 3 | 1095 | 239 | 153 |
| 4 | 673 | 147 | 78 |
| 5 | 415 | 82 | 57 |
| 6 | 323 | 51 | 39 |
| 7 | 194 | 35 | 32 |
| 8 | 154 | 25 | 23 |
| 9 | 97 | 17 | 17 |
| 10 | 82 | 18 | 21 |
| 11 | 52 | 10 | 13 |
| 12 | 44 | 4 | 12 |
| 13 | 18 | 1 | 9 |
| 14 | 15 | 2 | 8 |

† Genotypes were assigned to each strain for each SV call using Jasmine. Strains with the same genotype for a given SV call were considered to contain the same variant.

**Table S3:** Predicted consequences of deletions.

| Consequence type | Severity | Count |
|---|---|---|
| splice_donor_variant | HIGH | 733 |
| splice_acceptor_variant | HIGH | 724 |
| transcript_ablation | HIGH | 411 |
| frameshift_variant | HIGH | 223 |
| stop_lost | HIGH | 120 |
| start_lost | HIGH | 20 |
| stop_gained | HIGH | 2 |
| inframe_deletion | MODERATE | 91 |
| protein_altering_variant | MODERATE | 12 |
| splice_donor_5th_base_variant | LOW | 753 |
| splice_polypyrimidine_tract_variant | LOW | 124 |
| splice_region_variant | LOW | 69 |
| splice_donor_region_variant | LOW | 13 |
| stop_retained_variant | LOW | 7 |
| intron_variant | MODIFIER | 6064 |
| upstream_gene_variant | MODIFIER | 5110 |
| downstream_gene_variant | MODIFIER | 2034 |
| coding_sequence_variant | MODIFIER | 827 |
| 3_prime_UTR_variant | MODIFIER | 465 |
| 5_prime_UTR_variant | MODIFIER | 415 |
| non_coding_transcript_exon_variant | MODIFIER | 60 |

**Table S4:** Predicted consequences of duplications

| Consequence type | Severity | Count |
|---|---|---|
| transcript_amplification | HIGH | 401 |
| start_lost | HIGH | 1 |
| splice_polypyrimidine_tract_variant | LOW | 25 |
| intron_variant | MODIFIER | 1397 |
| upstream_gene_variant | MODIFIER | 730 |
| coding_sequence_variant | MODIFIER | 341 |
| downstream_gene_variant | MODIFIER | 106 |
| non_coding_transcript_exon_variant | MODIFIER | 87 |
| 3_prime_UTR_variant | MODIFIER | 34 |
| 5_prime_UTR_variant | MODIFIER | 15 |
| intergenic_variant | MODIFIER | 12 |

**Table S5:** Predicted Consequences of Inversions

| **Consequence type** | **Severity** | **Count** |
|---|---|---|
| start_lost | HIGH | 3 |
| splice_polypyrimidine_tract_variant | LOW | 2 |
| upstream_gene_variant | MODIFIER | 569 |
| coding_sequence_variant | MODIFIER | 514 |
| 5_prime_UTR_variant | MODIFIER | 267 |
| downstream_gene_variant | MODIFIER | 262 |
| 3_prime_UTR_variant | MODIFIER | 228 |
| intergenic_variant | MODIFIER | 20 |
| non_coding_transcript_exon_variant | MODIFIER | 18 |
| non_coding_transcript_variant | MODIFIER | 2 |

**Table S6:** Biological functions or phenotypes associated with environmental stimuli in overrepresented genes

| **Gene** | **Biological Function or Phenotype** | **SV Type** | **WormCat Terms[1]** | **Strains** |
|---|---|---|---|---|
| *srh-234* | Chemosensation[53] | DUP | C1_Transmembrane protein<br>C2_Transmembrane protein: seven transmembrane receptor<br>C3_Transmembrane protein: seven transmembrane receptor: srh | DL238<br>QX1794 |
| *str-208* | Missense mutations predicted to impair movement[55] | DEL | C1_Transmembrane protein<br>C2_Transmembrane protein: seven transmembrane receptor<br>C3_Transmembrane protein: seven transmembrane receptor: str | QX1794 |
| *hecw-1* | Inhibits *Pseudomonas aeruginosa* avoidance[59] | DEL | C1_Proteolysis proteasome<br>C2_Proteolysis proteasome: E3 | ECA396 |
| *clec-2* | Induced by *P. aeruginos*[60] | DEL | C1_Stress response<br>C2_Stress response: C-type Lectin<br>C3_Stress response: C-type Lectin | ECA36<br>ECA396<br>XZ1516 |
| *clec-10* | suppressed by *P. aeruginos*[60] | DEL | C1_Stress response<br>C2_Stress response: C-type Lectin<br>C3_Stress response: C-type Lectin | ECA396<br>JU310,<br>QX1794 |
| *clec-122* | Induced by *P. aeruginosa*[60] | DEL | C1_Stress response<br>C2_Stress response: C-type Lectin<br>C3_Stress response: C-type Lectin | ECA36 |
| *clec-143* | Induced by *P. aeruginosa*[60] | DEL | C1_Stress response<br>C2_Stress response: C-type Lectin<br>C3_Stress response: C-type Lectin | EG4725 |

| | | | | |
|---|---|---|---|---|
| *clec-174* | Induced by *P. aeruginosa*[60], *P. luminescens*, and *V. cholerae*[90] | DEL | C1_Stress response<br>C2_Stress response: C-type Lectin<br>C3_Stress response: C-type Lectin | DL238<br>XZ1516 |
| *clec-189* | May represent a novel immune effector (Pan et al., 2021) | DEL | C1_Stress response<br>C2_Stress response: C-type Lectin<br>C3_Stress response: C-type Lectin | EG4725<br>JU2526<br>QX1794<br>XZ1516 |
| | | DUP | C1_Stress response<br>C2_Stress response: C-type Lectin<br>C3_Stress response: C-type Lectin | EG4725<br>JU2526 |
| *cyp-35a4* | Inducible by caffeine[65] | DEL | C1_Stress response<br>C2_Stress response: detoxification<br>C3_Stress response: detoxification: CYP | XZ1516 |
| | | DUP | C1_Stress response<br>C2_Stress response: detoxification<br>C3_Stress response: detoxification: CYP | QX1794 |
| *cyp-33c5* | Cadmium detoxification[66] | DEL | C1_Stress response<br>C2_Stress response: detoxification<br>C3_Stress response: detoxification: CYP | EG4725<br>JU1400<br>JU2526 |
| *cyp-13a5* | Cadmium detoxification[66] | DEL | C1_Stress response<br>C2_Stress response: detoxification<br>C3_Stress response: detoxification: CYP | JU310<br>QX1794 |
| *cyp-33d3* | Inducible by Ethidium bromide[68]<br>Upregulated at low pH levels[67] | DEL | C1_Stress response<br>C2_Stress response: detoxification<br>C3_Stress response: detoxification: CYP | ECA396 |
| *cyp-33e2* | Inducible by caffeine[65] | DEL | C1_Stress response<br>C2_Stress response: detoxification<br>C3_Stress response: detoxification: CYP | ECA36<br>JU2526<br>XZ1516 |
| *ugt-14* | Inducible by albendazole, mebendazole, thiabendazole, and oxfendazole[70] | DEL | C1_Stress response<br>C2_Stress response: detoxification<br>C3_Stress response: detoxification: ugt | ECA36 |

1. Only terms with an adjusted *p*-value ≤ 0.05 were included.
